# Puromycin reveals a distinct conformation of neuronal ribosomes

**DOI:** 10.1101/2023.05.02.539124

**Authors:** Mina N. Anadolu, Jingyu Sun, Jewel T-Y. Li, Tyson E. Graber, Joaquin Ortega, Wayne S. Sossin

**Affiliations:** Department of Neurology and Neurosurgery, Montreal Neurological Institute, McGill University, Montreal, Quebec, H3A 2B4, Canada; Department of Anatomy and Cell Biology, McGill University, Montreal, Quebec H3A 0C7, Canada; Centre for Structural Biology, McGill University, Montreal, Quebec, H3G 0B1, Canada; Current affiliation: Children’s Hospital of Eastern Ontario Research Institute, Ottawa, Ontario, K1H 8L1, Canada

**Keywords:** Stalled polysomes, puromycin, neuronal RNA granule, Cryo-EM, ribosome

## Abstract

Puromycin is covalently added to the nascent chain of proteins by the peptidyl transferase activity of the ribosome and the dissociation of the puromycylated peptide typically follows this event. It was postulated that blocking the translocation of the ribosome with emetine could retain the puromycylated peptide on the ribosome, but evidence against this has recently been published (Hobson et al., 2020 https://doi.org/10.7554/eLife.60048; Enam et al., 2020 https://doi.org/10.7554/eLife.60303). In neurons, puromycylated nascent chains remain in the ribosome even in the absence of emetine, yet direct evidence for this has been lacking. Using biochemistry and cryo-electron microscopy, we show that the puromycylated peptides remain in the ribosome exit channel in the large subunit in a subset of neuronal ribosomes stalled in the hybrid state. These results validate previous experiments to localize stalled polysomes in neurons and provide insight into how neuronal ribosomes are stalled. Moreover, in these hybrid-state neuronal ribosomes, anisomycin, which usually blocks puromycylation, competes poorly with puromycin in the puromycylation reaction, allowing a simple assay to determine the proportion of nascent chains that are stalled in this state. In early hippocampal neuronal cultures, over 50% of all nascent peptides are found in these stalled polysomes. These results provide new insights into the stalling mechanisms of neuronal ribosomes and suggest that puromycylated peptides can be used to reveal subcellular sites of hybrid-state stalled ribosomes in neurons.

**Significance Statement:** Puromycin can be covalently linked to the nascent polypeptide chain on ribosomes, followed by dissociation of the puromycylated polypeptide. Here, we conclusively show that in stalled ribosomes isolated from neuronal RNA granules, the puromycylated peptide remains in the polypeptide exit tunnel of the ribosome. This validates previous data using this technique to localize stalled ribosomes in neurons and suggests a unique ribosomal conformation in these cells. Further evidence for the unique state of these ribosomes is the resistance of puromycylation to the inhibitor anisomycin, which prevents puromycylation in all other cellular contexts. These results provide insight into the mechanism underlying neuronal ribosome stalling and resolve a controversy about using puromycin to localize ribosome stalling in neurons.

## Introduction

Puromycin is a naturally occurring compound that mimics the 3’end of charged tRNAs. In the puromycylation reaction (1), the ribosome transfers the nascent polypeptide on the P-site tRNA to the amino group in the puromycin molecule in a reaction catalyzed by the peptidyl-transferase center (PTC). Since the labile ester bond found in charged tRNAs is replaced with a stable peptide bond in puromycin, the ribosome can no longer catalyze peptide joining reactions and translation stops. As the nascent polypeptide is no longer attached to a tRNA, the puromycylated nascent chain can diffuse away, presumably through the exit channel in the ribosome. It was postulated that prior to puromycylation, treatment of the translating ribosomes with elongation inhibitors, such as emetine, causes the nascent peptide to be retained in the ribosome. Therefore, this reaction, called ribopuromycylation (2), was used as a tool to label nascent polypeptides on ribosomes using anti-puromycin antibodies in order to detect the subcellular localization of actively translating ribosomes (3). However, recent data have provided evidence that even in the presence of emetine, puromycylated peptides still diffuse away from the ribosome (4, 5), bringing to question whether puromycylation methods in the literature are suitable for establishing subcellular localization of translation.

Local translation in polarized cells, such as neurons, requires the transport of translationally repressed mRNAs to regions like distal dendrites and synapses located far from the cell body. Translation needs to be reactivated later on at the appropriate time at these distal locations (6). One proposed mechanism is loading ribosomes onto mRNAs, pausing translation elongation and transporting stalled polysomes in neuronal RNA granules (7). Neuronal RNA granules were initially identified as large collections of ribosomes that sediment in sucrose gradients, the standard technique to separate monosomes and polysomes (8, 9). While some neurons contain dormant ribosomes (10) two recent studies examining ribosomes in RNA granules have shown that they are enriched in ribosomes with nascent chains that are stalled in the hybrid position with tRNAs in the A/P and P/E position (11, 12). Puromycin has been used to identify neuronal stalled polysomes in two ways. The first is independent of whether puromycylated peptides are retained on the ribosome. Run-off experiments using translation initiation inhibitors prevent the formation of new polysomes, thus depleting the population of actively elongating ribosomes and, consequently, puromycylation substrates. Given that stalled polysomes should resist run-off when initiation is blocked, an intact ribosome ostensibly permits puromycylation of nascent chains and their subsequent detection by immunoblotting with an antibody against puromycin (13). In cell lines with few stalled polysomes, these experiments reveal that run-off greatly decreases the number of puromycylated polypeptides. At the same time, there is only a small decrease in hippocampal cultures, suggesting a large proportion of nascent chains are on stalled polysomes (13). One can also do run-off experiments followed by ribopuromycylation to detect the location of stalled ribosomes in the cell. This second method depends on retaining the puromycylated peptide on ribosomes. These experiments show stalled ribosomes in puncta in neurites that colocalize with markers of RNA granules, suggesting that RNA granules contain abundant stalled ribosomes (13). However, if emetine does not block the dissociation of the puromycylated peptides, the puncta observed could be puromycylated polypeptides dissociated from ribosomes that are still present in the liquid-liquid phase-separated RNA granules (5).

An alternative method for detecting nascent polypeptides is the SunTag technique, in which the translation of a repeated peptide epitope can be detected in real time by co-transfection of a GFP-tagged single chain antibody that recognizes this epitope (14). During translation, this leads to the localization of many molecules of GFP forming a punctum that marks the location of polysomes. Adding a degron (a sequence that leads to protein degradation) at the end of the translated peptide leads to the loss of puncta when translation is terminated due to the diffusion and degradation of the protein after translation (15). Indeed, in non-neuronal cells or neuronal soma, the addition of puromycin leads to the loss of SunTag puncta, even in the presence of emetine (5, 16) suggesting that nascent polypeptides are released. In contrast, neurites exhibit SunTag puncta resistant to puromycin (16). Importantly, these puncta colocalize with markers of RNA granules (16). It is still unclear whether the polypeptides recognized by the GFP tagged single chain antibody in RNA granules after puromycin treatment are dissociated from the ribosomes but protected from degradation because of the liquid-liquid phase-separated nature of the RNA granules (5).

Our study directly tests whether neuronal ribosomes retain puromycylated nascent chains using biochemical fractionation experiments and cryo-electron microscopy (cryo-EM). Our experiments demonstrate that the puromycylated polypeptide remains associated with the stalled ribosomes in neuronal RNA granules after puromycin treatment. Imaging these ribosomes using cryo-EM showed a population containing the nascent chain retained inside the exit channel with a puromycin molecule attached to the C-terminal end of the polypeptide at the P-site of the 60S subunit. Additional experiments also support a distinct conformation of these ribosomes. Anisomycin binds to the A site of the ribosome, competes with puromycin and thus prevents puromycylation. However, puromycylation is only partly blocked by anisomycin in ribosomes from neuronal RNA granules. The altered ability of anisomycin to compete with puromycin provides a convenient assay to determine the approximate percentage of neuronal ribosomes in this conformation. We discuss the implications of these observations in understanding the mechanisms of stalling in neuronal ribosomes.

## Results

### Puromycylated nascent peptides in RNA granules segregate with ribosomal, not soluble fractions

RNA granules enriched in stalled ribosomes can be found in the pellet of sucrose gradients normally used to separate monosomes from polysomes in cellular lysates (9, 17–20). We enriched for RNA granules from whole brain lysates of five day old rats of both sexes using a short centrifugation protocol (17, 19) and then treated the resuspended granule fraction (GF) with the same concentration of puromycin (100 μM) used in previous ribopuromycylation experiments (13) (Figure 1A). No emetine was used since emetine does not affect ribopuromycylation of stalled ribosomes in neurons (16). Resedimenting the granule fraction by centrifugation after puromycylation indicated that a large percentage of the puromycylated nascent chains were not released into the supernatant by this treatment but instead sedimented with the RNA granules (87%+3% of the puromycin-specific immunoreactivity pelleted, n=3; Figure 1B). To determine whether the puromycylated nascent peptides were retained on the ribosomes in the RNA granule or in the liquid-liquid phase of RNA granules, we dissociated the polysomes in the granule into monosomes by treating them with nuclease (17). The monosomes were then purified by an additional sucrose gradient sedimentation (Figure 1C). Few soluble proteins were expected from this preparation since the starting material was the pellet. However, because the protein used to inactivate the nuclease (SuperasIN) cross-reacts with our antibody against ribosomal protein eS6 (Figure 1D; Supplemental Figure 1A), we were able to use this protein as a marker for soluble proteins in this fractionation. Monosomes, initially identified by eS6 immunoreactivity and later by EM (see below), were concentrated in fractions 2 and 3 of this sucrose gradient (17)(Figure 1D; Supplemental Figure 1B). Most puromycylated peptides were found in the same fractions as the monosomes, with only minor amounts found in fraction 1 (the soluble fraction) (Figure 1D; Supplemental Figure 1B). Cleavage of the polysomes in the GF was incomplete (17) thus, some of the ribosomes (anti-eS6 staining) and puromycylated nascent chains (anti-puromycin) resedimented in the pellet (Figure 1D, Supplemental Figure 1B). Performing the puromycylation reaction at 37 °C or monitoring large ribosomal subunits instead of small subunits gave similar results (Supplemental Figure 1C). This data suggests that the puromycylated peptides segregate with and are retained on ribosomes in granules derived from whole brain homogenates rather than diffusing away through the ribosome exit channel.

**Figure 1.**
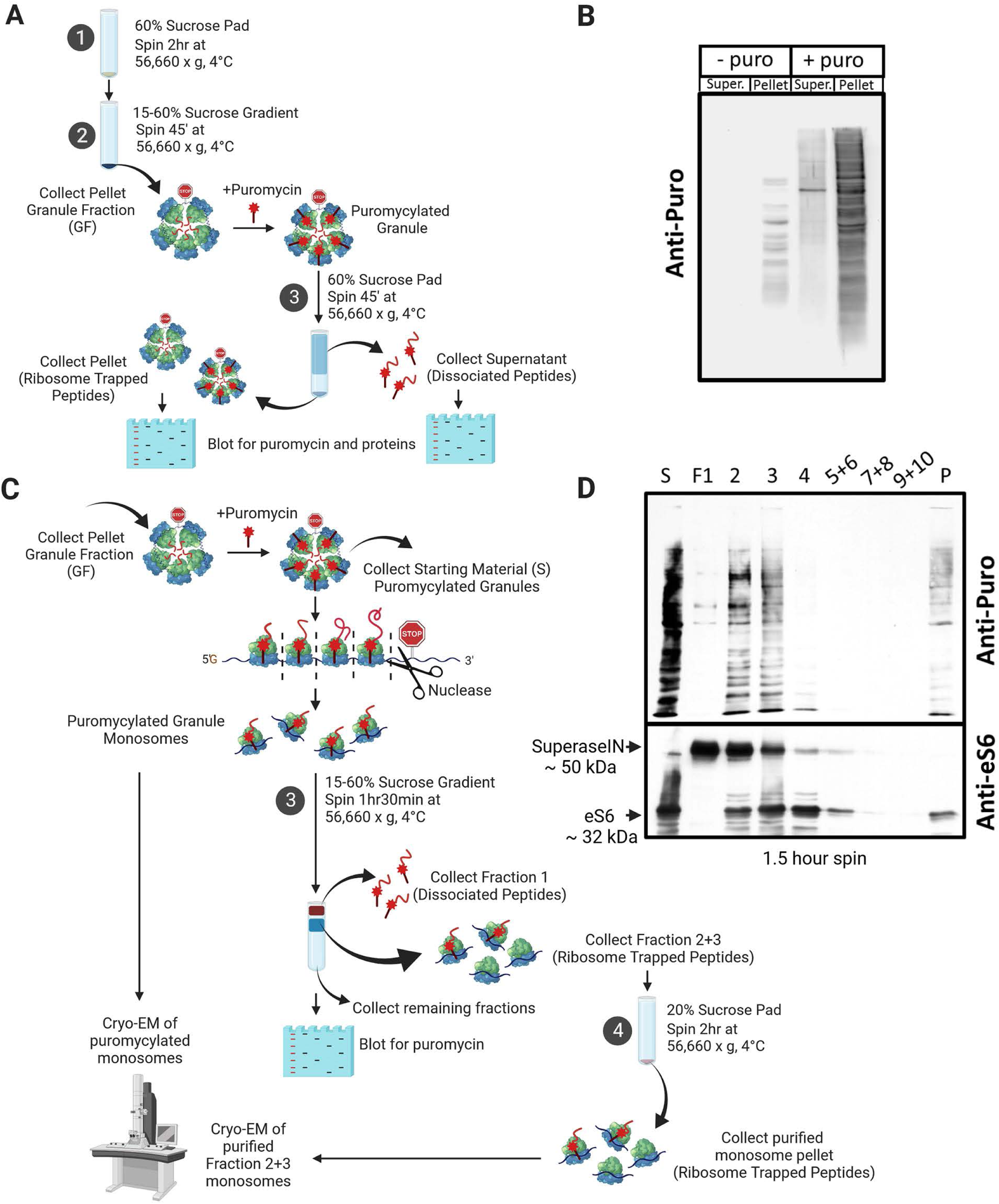
Puromycylated peptide are retained by ribosomes in the granule fraction. A) Outline of how granule pellets are purified from P5 whole brain lysate and puromyclated, followed by centrifugation to observe whether puromycylated nascent peptides escape the ribosomes, using immunoblotting. B) Immunoblot with anti-puromycin to determine if puromycylated nascent chains resediment. C) Outline of how granule pellets from P5 whole brain lysate are puromycylated and treated with nuclease to produce monosomes to observe whether puromycylated nascent peptides stay with the monosomes (Fraction 2+3) or escape to the soluble fraction (Fraction 1) using immunoblotting and cryo-EM. Samples treated with nuclease were run on the sucrose gradient for 1 hour 30 minutes to ensure proper separation of monosomes (F2) from the soluble fraction (F1). D) Immunoblot of starting material (puromycylated GF) and sucrose gradient fractions after nuclease digestion, immunostained against puromycin and S6 ribosomal protein; S: start, F1: soluble fraction, F2: monosomal fraction, P: pellet. Fractions 5 and 6, 7 and 8, 9 and 10 were pooled to fit all fractions into the gel. A band of ∼50 kDa size was observed when immunostaining with the anti-S6 antibody, which was identified to be caused by SuperaseIN used to quench the RNase I activity (See Supplemental Figure 1).

### Cryo-EM of puromycylated ribosomes reveals a novel class

To further investigate whether puromycylated nascent peptides remained in the ribosome, we examined the monosomes from fractions 2 and 3 using cryo-EM. We have previously used this method to analyze the monosomes after nuclease treatment of the pellet and found a large fraction of ribosomes were in the hybrid state (17), consistent with other studies of ribosomes in neuronal granules (12). We reclassified the dataset from Anadolu et al., 2023 to include dissociated 60S subunits since these were discarded in the initial analysis, but may represent an important class after puromycin treatment if, as in other ribosomes, puromycin causes dissociation of the nascent chain and the dissociation of the ribosomal subunits (4). In reclassifying the non-puromycylated ribosomes, class 1 represented 75% of the population and exhibited tRNA molecules in hybrid A/P and P/E states. Class 2 was 12% of the ribosomes and contained a tRNA in the P-site. The remaining particles were dissociated 60S subunits, accounting for 13% of the population (Figures 2 & 3A).

**Figure 2.**
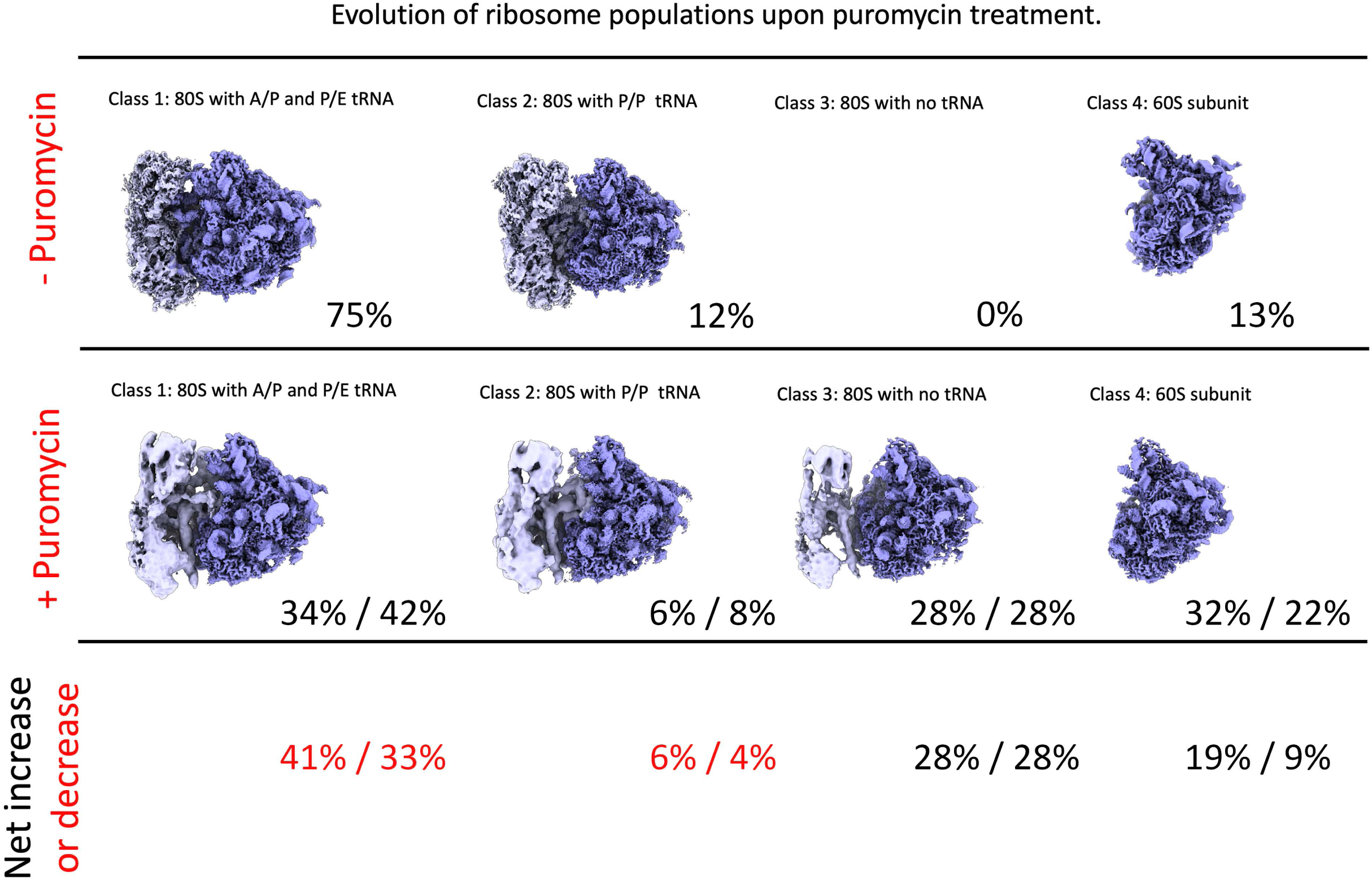
Evolution of the ribosome populations upon puromycin treatment. The cryo-EM particle datasets obtained for the 80S ribosomes in the granule fraction of the sucrose gradient after nuclease treatment and the equivalent dataset obtained for the same sample with an additional puromycin treatment were subjected to image classification approaches (Supplementary Figure 2 & 3) and Methods). The table shows the various classes found in each dataset and the percentage of the total population that each class represented. The last row of the table shows the net increase (black numbers) or decrease (red numbers) for each class. The two percentage numbers in the ‘+ puromycin’ and ‘net increase or decrease line’ refer to the classification results when the sample that was analyzed by cryo-EM was processed as described in Figure 1A & 1C (first set of percentages) or according to our previous analysis (17), in which after treating the granule fraction with puromycin and nuclease, samples were directly applied to cryo-EM grids.

**Figure 3.**
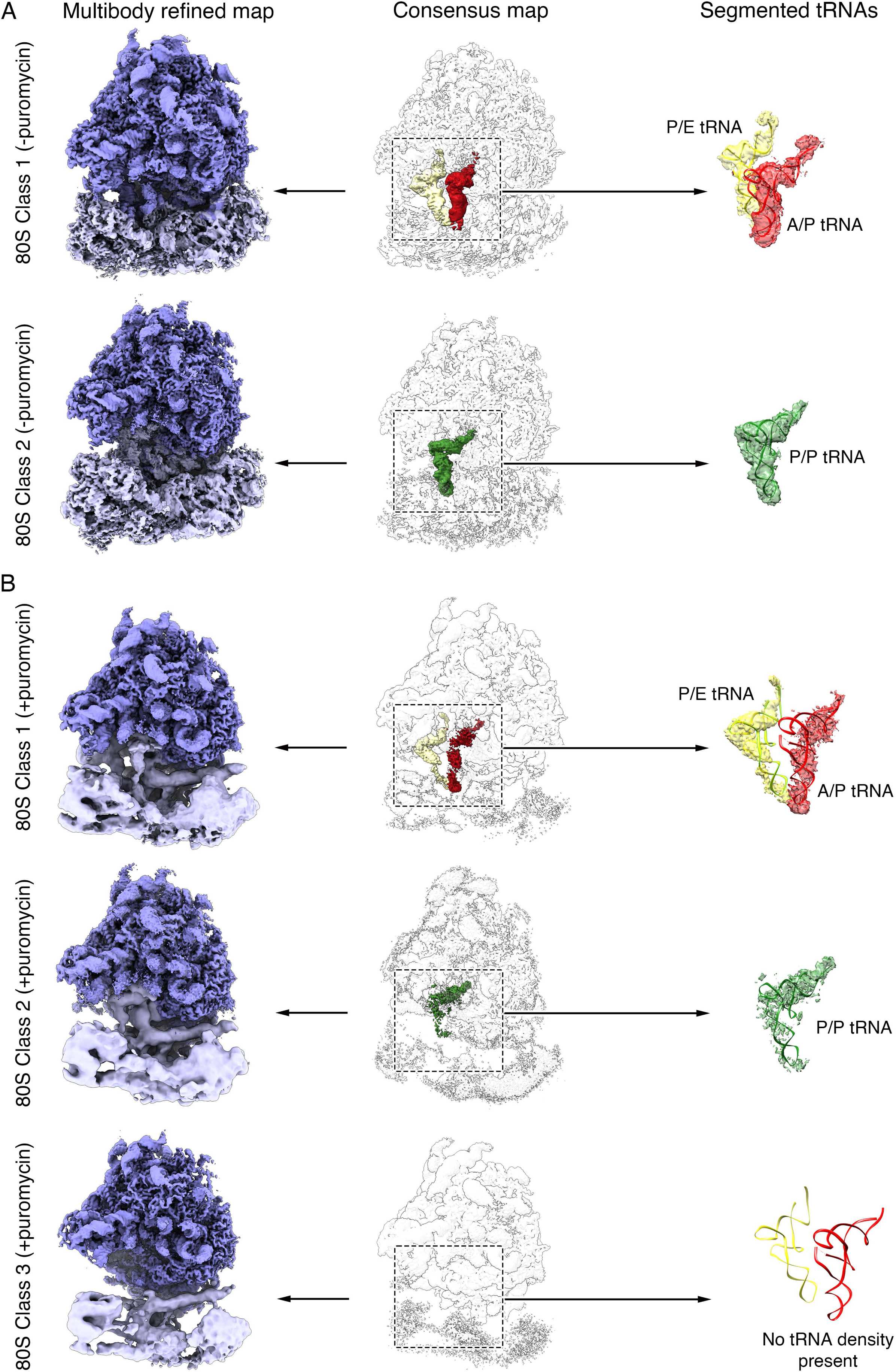
Cryo-EM analysis of puromycin-treated neuronal ribosomes from the granule fraction. (A) Previously published (non-puromycylated) cryo-EM maps of the two classes of 80S ribosomes found in the granule fraction of the sucrose gradient after nuclease digestion. The left panels show a side view of the cryo-EM maps of the 80S ribosomes (from multi-body refinement) with the 40S subunit and 60S subunit colored in cyan and dark blue, respectively. The middle panels show the cryo-EM maps of the 80S ribosomes (consensus maps) as transparent densities to show the tRNA molecules present (or absent) in these classes. The right panels show the tRNA molecule densities segmented out of the maps. Images of the cryo-EM map were prepared from EMDB entries 29538 (class 1, consensus), 20726 (class 1, 40S multi-body refinement), 28727 (class 1, 60S multi-body refinement), 29539 (class 2, consensus), 26517 (class 2, 40S multi-body refinement), 26518 (class 2, 60S multi-body refinement). The ribbon representation of the A/P and P/E tRNA molecules docked into the densities in the right panels was obtained from PDB ID 6HCJ. The P/P tRNA was obtained from PDB ID 6R5Q. (B) Classes of 80S ribosomes found in the granule fraction of the sucrose gradient after puromycin treatment and nuclease digestion. Panels on the left, middle and right are arranged as in panel A.

Image processing approaches of the cryo-EM images (see Methods; Supplemental Figures 2 and 3) from the monosomes after puromycin treatment revealed two classes similar to class 1 and 2 in the untreated dataset. The percentage of particles that classes 1 and 2 represented decreased by 41% and 6%, respectively, relative to the untreated sample (Figure 2). In addition, we identified a novel class of 80S ribosomes in the puromycin-treated sample, which we called class 3 (Figure 2). This class represented 28% of the population and showed no tRNA density in the interface between the two subunits (Figure 3). We also noticed a 19% increase in free 60S subunits, now called class 4, in the puromycin-treated sample (Figure 2). To ensure that changes in the classes and the new class 3 structure were not due to the additional centrifugation steps to purify and concentrate the monosomes following puromycylation and nuclease treatment (Figure 1C), we also analyzed monosomes using cryo-EM after direct sedimentation of the puromycin and nuclease treated granule fraction, as performed in Anadolu et al., 2023. We found the distribution of the four classes of ribosomes (class 1: 42%, class 2: 8%, class 3: 28% and class 4: 22%) comparable to the sample with additional centrifugation steps (Figure 2). These results demonstrated that these changes were not due to the additional time and centrifugation used to purify and concentrate monosomes.

Classes 1 and 2 of the puromycin-treated preparation (Figure 3) resembled classes 1 and 2 in the untreated sample and contained the same type of tRNA molecules (Figure 3). However, the density for the 40S subunit was highly fragmented in all the puromycin-treated samples and did not show high-resolution features (Figure 3) even after applying multibody refinement. The 40S subunit in class 3 also showed severe fragmentation. In contrast, the maps obtained for the large subunits in all puromycin-treated classes refined to a resolution of ∼ 3Å (Supplemental Figure 4). The densities representing the A/P tRNA in class 1 and P/P tRNA in class 2 were also partially fragmented in the puromycin-treated sample (Figure 3) but showed complete densities in the untreated monosomes (Figure 3).

To rule out that misclassification was not the cause of the observed resolution degradation and fragmentation of the 40S density, we analyzed the 2D class averages obtained from the particles in each class. We found that the 2D class averages for classes 1 and 2 of non-treated preparation showed high-resolution features in the 60S and 40S subunits (Supplementary Figure 5A). However, class averages derived from the three classes in the puromycin-treated preparation showed blurred densities for the 40S subunits, but more importantly, they did not show an absence of this density (Supplementary Figure 5B). These results suggested that the resolution degradation and fragmentation of the 40S density was caused by their enhanced mobility in the puromycin-treated ribosomal particles.

Consequently, we used 3D variability analysis to explore the enhanced movement of the 40S subunit in classes 1, 2 and 3 in the puromycin-treated sample, and we compared it to classes 1 and 2 in the untreated sample. Particles from each class were analyzed separately, and the resolved variability components showed mainly three type of motions and the range of these motions for each class. The first component of the 3D variability analysis identified a detachment of the bottom part of the 40S subunit from the 60S subunit. This motion was present in all classes, but the range of this motion was much more prominent in the three classes from the puromycin-treated sample (videos 4, 7 and 10) than in the classes from the untreated sample (Videos 1 and 2). The motion described by the second component involved the ratchet-like inter-subunit movement in which the 40S subunit rotates with respect to the 60S subunit. Class 1 from the untreated sample lacked this motion. Still, it was observed in all other classes, and it was again more prominent in the classes from the puromycin-treated sample (Compare Video 3 (class 2, untreated) with Videos 5, 8 and 11(classes 1, 2 and 3, puromycin treated)). Finally, the three classes from the puromycin-treated sample showed an additional type of motion that involved sliding the 40S subunit over the 60S subunit (Videos 6, 9 and 12). This motion was not observed in class 1 and 2 in the untreated samples. A pattern observed in the ratchet-like inter-subunit movement in the classes from the puromycin-treated sample was that the degree of motion increased from the class with an A/P and P/E tRNAs (class 1, Videos 5) to the class exhibiting only a P/P tRNA (class 2, Videos 8) and it was even more significant in the class that did not contain a tRNA molecule (class 3, Video 11).

These results suggested that the tRNA molecules and 40S subunit in the treated sample are not as tightly bound and adopt a more flexible conformation, implying that puromycin treatment destabilizes these two ribosome components. Similar fragmentation and reduced resolution were seen in the puromycylated and nuclease treated monosomes directly deposited on grids, and thus, these differences from our previous results are not due to the additional centrifugation steps required to purify and sediment monosomes.

### Puromycin treatment caused formation of puromycylated peptides that are retained in the stalled ribosomes

Next, we focused our analysis on the A and P sites of the large subunit and their connection to the nascent peptide, where we might expect to find evidence of densities related to puromycin. All 3’ends of tRNAs terminate with a cytosine, cytosine, adenosine (CCA) sequence. Critically, puromycin mimics the 3’ adenosine of a tRNA charged with a modified tyrosine (Figure 4A) and is a substrate of the peptidyl-transferase reaction (1, 21). In our recently published structures of class 1 80S ribosomes from the granule fraction (17), the densities for the 3’ CCA nucleotides in the A/P tRNA are visible. In addition, the terminal adenosine is, as expected, covalently linked to the C-terminal end of the nascent polypeptide chain extending from the P site into the exit channel of the ribosome (Figure 4B and Supplemental Figure 6; top middle panel). In the new class 3 ribosome identified in the puromycin-treated sample and without a tRNA molecule in the interface region, we observed a distinct density attached to the C-terminal end of the nascent polypeptide of the size and shape of the puromycin molecule (Figure 4C, bottom panel). Therefore, this structure provides visual evidence that the nascent polypeptide in this ribosome population was transferred to the puromycin molecule. Moreover, in these structures, the puromycylated peptide is retained in the exit channel and does not diffuse away from the ribosomes as described for non-neuronal ribosomes (Supplemental Figure 6, bottom middle panel) (4, 5).

**Figure 4.**
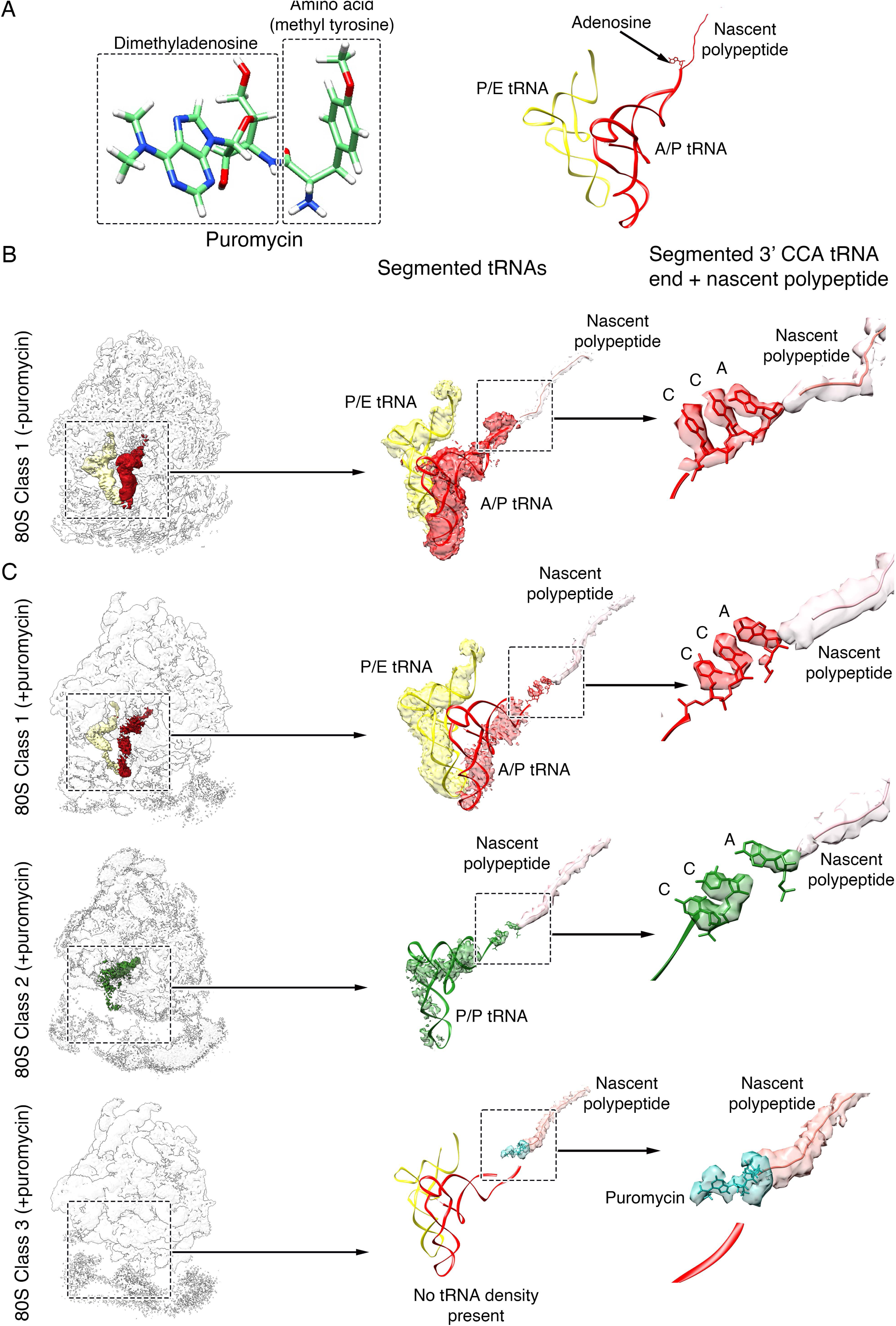
Cryo-EM analysis of densities proximal to the conserved CAA nucleotides at the 3’ end of tRNA molecules in neuronal ribosomes from the GF. (A) Comparison of the 3D structure of puromycin (left) with the 3’ terminus of the tRNA molecule bound to the nascent polypeptide (right). Comparison of the 80S ribosome class 1 existing in the granule fraction of the sucrose gradient after nuclease digestion (B) with the three classes of 80S ribosomes found in the same sample after puromycin treatment (C). The left panels show the consensus cryo-EM maps as transparent densities to show the tRNA molecules present in these classes. The middle panels show the tRNA molecule and nascent polypeptide chain densities segmented from the maps. The right panels show a zoomed-in view of the 3’end of the A/P and P/P tRNA molecule connected to the polypeptide chain. In the case of the puromycin treated 80S class 3, no tRNA molecules are observed in the cryo-EM map. Instead, a density of the size and shape of the puromycin molecule is observed attached to the nascent polypeptide chain. Images on panel B were prepared from EMDB entries 29538 (consensus map).

In addition, we analyzed the densities representing the 3’ CCA nucleotides of the A/P and P/P tRNA molecules in the cryo-EM maps obtained for classes 1 and 2 in the puromycin treated samples (Figure 4C, top and middle panel). Consistent with the fragmented appearance of the entire tRNA molecule in these cryo-EM structures, the density representing the CCA 3’ end nucleotides of the tRNA molecule was not as well defined when compared to that of the equivalent class in the untreated samples. In particular, the density assigned to adenosine was detached from the density assigned to the CC at 3’ end nucleotides of the tRNA but connected to the nascent polypeptide that was visible in the exit channel (Figure 4C, top and middle panel and Supplemental Figure 6, top and bottom left panels). However, given the resolution of the features in this region of the map, it was not possible to discriminate whether this density corresponded to puromycin or to the 3’ adenosine of the tRNA molecule that was present in a flexible conformation causing density fragmentation. While inconclusive, these results are consistent with puromycylation having occurred in these samples, followed by an increase in the mobility of the unconnected tRNA. This may represent an intermediate state between the start of the puromycylation reaction and the loss of the tRNAs, as observed in class 3. The free 60S subunits in the sample treated with puromycin (class 4) lacked a nascent polypeptide in the 60S exit channel (Supplemental Figure 6, bottom right panel), and this result is consistent with the dissociation of the nascent chain and separation of the two subunits as is seen in non-neuronal puromycylated ribosomes. However, there are free 60S subunits present even before puromycylation, and we did not detect free 60S subunits (as monitored by L7 staining) or free 40S subunits (as monitored by eS6 staining) migrating in the gradient without nuclease treatment (Supplemental Figure 7), suggesting association of free 60S subunits in the ribosome clusters found in the granule fraction (17) and continued association of both subunits after puromycylation treatment.

### Anisomycin competes poorly with Puromycin in neuronal ribosomes

Overlapping previously obtained structures of the ribosome in complex with anisomycin (22, 23) and puromycin (23, 24), we found that the anisomycin site always partially overlaps with the puromycin binding site. Anisomycin binds in the groove that accommodates the charged amino acids attached to tRNAs and thus prevents charged tRNAs from binding in the A site (23). Puromycin binds lower on the A site, mimicking the 3’end of aminoacyl tRNA with the tyrosine ring overlapping the anisomycin binding site (23). Consequently, anisomycin can compete with puromycin in the puromycylation reaction (2). However, we found that in neuronal ribosomes from the GF, considerable puromycylation occurs in the presence of anisomycin (Figure 5A). This peculiar resistance of the puromycylation reaction to anisomycin should allow us to assess what percentage of ribosomes in a given cellular context are in the same state as the ribosomes in the GF. To this end, we examined hippocampal cultures, cortical cultures and HEK293 cells for anisomycin-puromycin competition (Figure 5B). As expected, anisomycin completely blocked puromycylation in HEK cells, in which few ribosomes are stalled (13). Instead, puromycylation was resistant to anisomycin in cultured hippocampal neurons, similar to the ribosomes in the GF (Figure 5B, quantified in Figure. 5C), in which over 80% of the ribosomes with nascent peptides are in the hybrid state (class 1; Figure 2). Surprisingly, anisomycin competes significantly better in cortical cultures suggesting that fewer ribosomes in cortical cultures are in this state than in hippocampal cultures (Figure 5B, quantified in Figure 5C). This finding indicates that the binding site of the two antibiotics in neuronal ribosomes may not overlap when they are stalled at the hybrid state, meaning that ribosomes in neurons uniquely respond to treatments with puromycin and anisomycin.

**Figure 5.**
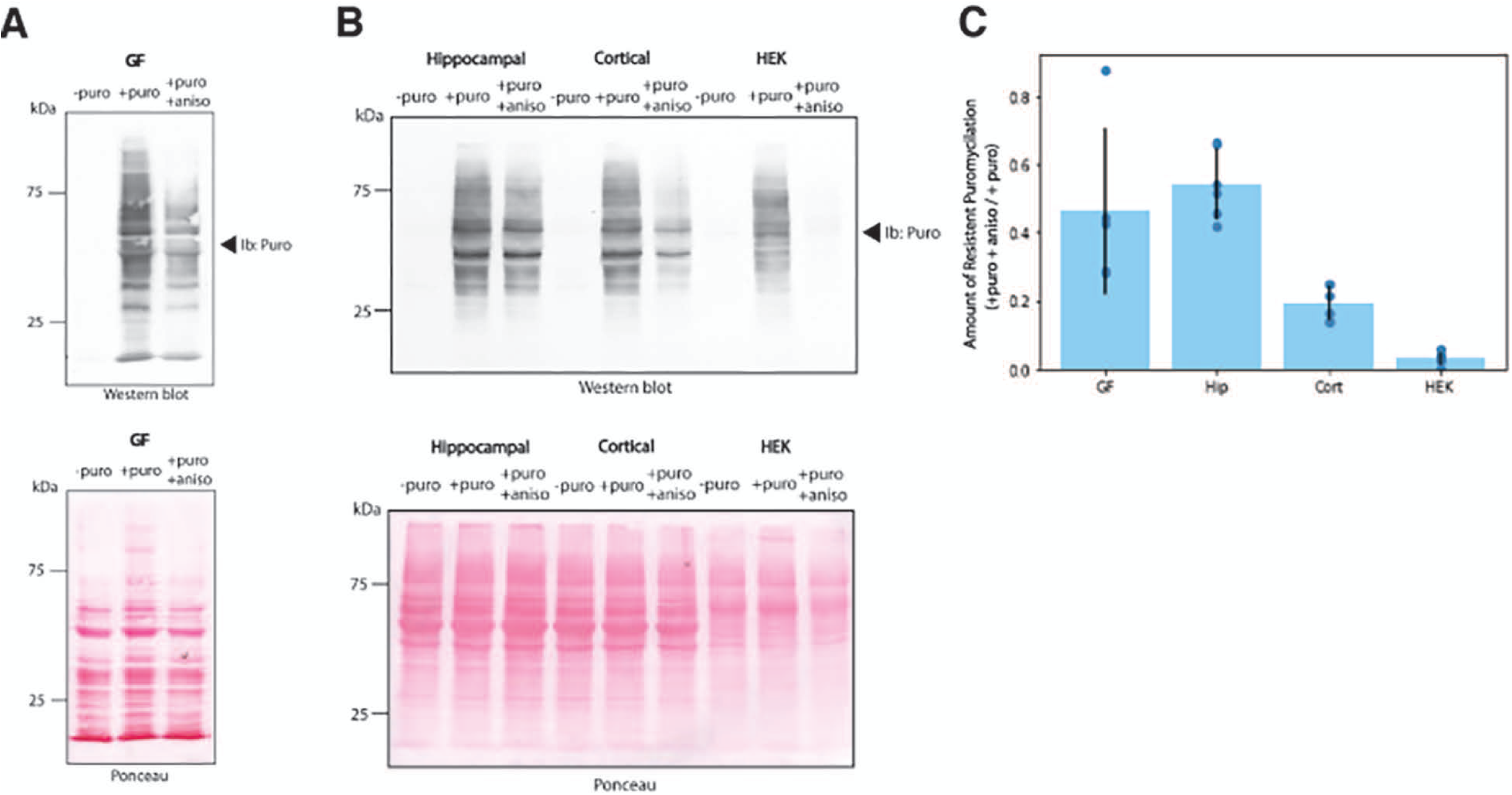
Competition of the puromycylation reaction with anisomycin. (A) Example of inhibition of puromycylation (100 µM) by anisomycin (100 µM) in the GF. The top panel shows the immunoblots developed with the ani-puromycin antibody. The bottom panels are the same membranes stained with Ponceau before immunoblotting. B) Example of inhibition of puromycylation (100 uM) by anisomycin (100 µM) in live cultures of hippocampal neurons, cortical neurons or HEK293 cells. Immunoblots and Ponceau stained blot are arranged as in A. (C) Quantification of the amount of puromycylation resistant to anisomycin treatment. ANOVA F(19,3)=20.9, p<0.0001; Post-Hoc tests; all groups are significantly different from each other (p<0.01) except for Pellet vs Hippocampal neurons (P>0.05).

## Discussion

The Class 3 cryo-EM structure obtained from puromycylated RNA granules, harbouring a puromycin density adjacent to a nascent peptide chain retained on the exit channel of the large ribosomal subunit (Figure 4 and Supplementary Figure 6), strongly demonstrates the retention of puromycylated peptides in at least a subset of neuronal ribosomes. This observation is supported by previous experiments in neurons using ribopuromycylation, in which immuno-labelled puromycin localizes to the same puncta as the ribosomal proteins and granule-associated proteins (13, 25), and using Sun-Tag where puromycin did not cause loss of the Sun-Tag puncta (16). The ability to detect a retained puromycylated peptide was questioned in the past due to the anticipated difficulty for the anti-puromycin antibody to access its epitope at the entrance of the exit channel (5). However, in this argument, researchers assumed that the tRNA molecules in this structure were retained, and these contributed to the steric blockages of the antibody. Our findings of increased mobility of the 40S subunit in all 80S samples treated with puromycin (Fig. 3B and Videos 4-12) may allow access of the anti-puromycin antibody to its epitope even in the presence of tRNAs. Together these data conclude that ribopuromycylation studies performed in neurons (13, 25, 26) correctly localize stalled ribosomes because the puromycylated nascent peptides do not dissociate from these ribosomes.

Puromycylation of the hybrid position requires that puromycin has access to its binding site in the presence of the A/P site tRNA. Although tRNAs cannot access the A site of the large subunit in this ribosomal conformation, puromycin which mimics only the 3’ end of tRNA can do so. However, puromycylation in the hybrid state is much slower than when the A site is empty (27–30). While this is interpreted to be because of decreased efficiency of the ribosome in catalyzing peptide joining due to the altered position of the acceptor amino acid (28), it is possible that a difference in the way puromycin binds in the hybrid state ribosome could explain the slow speed of puromycylation. A difference in how puromycin binds to the hybrid state ribosome is also consistent with the lack of competition between anisomycin and puromycin in neuronal ribosomes shown in our data. However, even ribosomes slowly cycling between hybrid states and classical states would become puromycylated as soon as they cycled, so neuronal stalled ribosomes would have to be locked in the hybrid state. Alternatively, it is possible that some altered structure of the stalled neuronal ribosome, not present in all hybrid state ribosomes, may contain an altered puromycin binding site and consequently show a decreased competition between anisomycin and puromycin. Further experiments are required to differentiate these two models.

Consistent with ribopuromycylation in neurons being specific to the hybrid state stalled ribosomes, we have noticed that a large concentration of puromycin is required to observe ribopuromycylated puncta in neurites (Supplemental Figure 8). It should be noted that most usage of puromycin in labeling protein synthesis uses a low concentration of puromycin (1-5 μM) (3) and thus may not capture the nascent chains in hybrid state stalled ribosomes.

### Implications for stalling mechanism

While our data show that the puromycylated peptide remains in the peptide exit channel in the ribosome (Supplemental Figure 6), it does not explain the retention mechanism. One possibility is that the peptide was already stalled in the exit channel before puromycylation occurred and that peptide stalling is a mechanism for neurons to stall polysomes. This stalling is unlikely to be related to issues, such as polyproline stretches, where stalling is due to the slowing of peptide joining and thus, the hybrid state is not populated, allowing eIF5A to occupy the site of the E site tRNA (31, 32). In contrast, stalling induced by a drug that binds to a pocket in the peptide exit channel stalls translation in the hybrid state (33), as does stalling of the SecM peptide in bacteria (34). The large proportion of ribosomes in neuronal RNA granules that are in the hybrid state seen in ours and others data (12) is far greater than seen in other cryo-EM studies of ribosomes from non-neuronal cells which range between 15-30% (35–38). Stalling mediated by the XBP1 peptide is mainly in the non-rotated state, although a fraction of the structures is in the hybrid state (39). In the cryo-EM structure calculated for class 3 from the sample treated with puromycin, we analyzed whether the puromycin induced a kink in the nascent chain. However, we did not observe any clear difference in the conformation of the nascent peptide or in the PTC with the structure previously obtained for the hybrid state ribosomes (class 1) in the untreated sample (17) (Supplementary Figure 6). Codons encoding acidic amino acids (Asp and Glu) are close to two-fold enriched in peaks of ribosome-protected mRNA fragments recovered from the granule fraction (17). This finding is consistent with a stalling mechanism where a neuronal-specific factor facilitates stalling of peptides rich in acidic residues. Acidic amino acids are linked to faster elongation in yeast (40). Aspartic acid is enriched at the A site in developing neocortex suggesting slow elongation with this residue in the A site (41). However, slowing of elongation by a specific amino acid in the A site suggests slow peptide joining and as discussed above, this is not consistent with stalling in the hybrid state. Other factors potentially contributing to other stalling mechanism, such as EBP1 which binds to the exit channel of neuronal ribosomes during early development (42) are unlikely to play a role. EBP1 is found bound to both hybrid state and non-hybrid state ribosomes (42) and thus, is unlikely to be a critical factor in the stalling mechanism proposed here. We did not observe this density in our cryo-EM structures.

We also noted that there are cryo-EM densities connecting the stalled nascent polypeptide with nucleotides surrounding the exit channel (Supplemental Figure 6). Therefore, a second possibility to explain the lack of release of the puromycylated peptide is by a mechanism that combines the effect of still uncharacterized interactions of the puromycin molecule in the PTC and the nascent peptide with surrounding nucleotides of the 28S rRNA in these two areas. Under this scenario, the puromycin and the nascent peptide may be retained as a direct consequence of the stalling mechanism, which remains uncharacterized.

### Implications for neuronal development

Our results indicate that stalled polysomes constitute a large proportion of ribosomes during this stage of neuronal development (P5 brains). Indeed, in cultured hippocampal neurons, a major model for examining neuronal translational control (43), most nascent peptides are retained in stalled polysomes based on both the reduced competition with anisomycin (Figure 5) and previous results showing that puromycylation is not significantly reduced by initiation inhibitors (13). Since the most abundant mRNAs in these stalled polysomes encode cytoskeletal proteins such as tubulin, actin, microtubule-associated proteins and motors (17), it suggests that this mechanism is critical for the growth of axons and dendrites at this developmental stage. However, this is difficult to test, because without a known stalling mechanism, it is impossible to specifically block the formation of stalled ribosomes. Later in development, and in other neuronal cell types, the percentage of stalled ribosomes may decrease. Thus, other forms of local transport and translation, such as mRNAs stalled at initiation may play a more significant role in the translational landscape later in development. Nevertheless, we have shown that mGLUR-LTD in mature neuronal cultures and slices is mainly mediated by the reactivation of stalled polysomes (13, 25), so this mode of translational control persists in the adult.

### Conclusions

In neurons, puromycylation of a subset of ribosomes does not lead to the release of the puromycylated peptide from the exit channel and dissociation of the ribosomal subunits as would be expected (4, 5). This suggests the possibility that stalling of these ribosomes is due to a stall of the nascent peptide in the exit channel. We found that in these ribosomes, there is a decrease in the ability of anisomycin to block the puromycylation reaction, providing an assay to examine the proportion of ribosomes that occupy this state. In growing hippocampal neurons, a large proportion of nascent chains on ribosomes, presumably over 50% are stalled in this state. Surprisingly, cortical neurons have a smaller proportion of these ribosomes. Our results suggest that understanding how neuronal ribosomes are stalled and why their proportions differ between neuronal populations will be critical in understanding the role that this form of regulated translation plays in neuronal function and neurodevelopment.

## MATERIAL AND METHODS

### Purification of the RNA Granule-Enriched Fraction

RNA granules were enriched from whole brain homogenate harvested from five-day-old (P5) Sprague Dawley rats of both sexes (Charles River Laboratories), as previously described (17). Five P5 rat brains were homogenized using a Caframo Mixer [Cole Parmer Canada P1086085] in RNA granule buffer (20 mM Tris-HCl pH 7.4 [Fisher; BP152-1], 150 mM NaCl [Fisher; BP358-212], 2.5 mM MgCl_2_ [Fisher; M33-500]) supplemented with 1 mM DTT [Sigma; D9163], 1 mM EGTA [Sigma; E8145], EDTA-free protease inhibitor [Roche; 04693132001]. Homogenate was centrifuged 15 minutes in a Thermo Scientific T865 fixed angle rotor at 6117 x g at 4°C to spin down cellular debris. The supernatant was treated with 1% IGEPAL CA-630 [Sigma; I8896] for 5 minutes at 4°C on a rocker. The sample was then loaded onto a 2 ml 60% sucrose [Calbiochem; 8550] cushion (dissolved in supplemented RNA granule buffer) in a Sorvall 36 ml tube [Kendro; 3141, Thermo Scientific], filled to top with additional RNA granule buffer and centrifuged for 2 hours in a Thermo Scientific AH-629 swing-bucket rotor at 56660 x *g* at 4°C to achieve the polysome pellet. The polysome pellet was re-suspended in RNA granule buffer, gently dounced and loaded over a 15-60% (w/v) linear sucrose gradient (made with RNA granule buffer) that was prepared in advance using a gradient maker [Biocomp Gradient Master] and centrifuged for 45 minutes at 56660 x *g* at 4°C in an AH-629 swing bucket rotor. The pellet from this centrifugation was rinsed once with RNA granule buffer and then resuspended with 1ml of the same buffer. The resuspended pellet enriched in RNA granules is called the granule fraction (GF). See Puromycylation and Nuclease Treatment sections below for how the GF was treated (Figure 1).

### Puromycylation of Granule Fraction

The resuspended GF was split evenly into two 1.5ml microtubes and incubated with 100 µM puromycin (Sigma Aldrich P7255) or vehicle in RNA granule buffer for 5 minutes at 4°C or 37^0^C. To see whether the puromycylated peptide escapes the ribosomes in the GF, the puromycylated GF and vehicle treated control GF were loaded onto 60% sucrose pads in open top polycarbonate centrifuge tubes (Beckman Coulter #343776) filled to top with RNA granule buffer and centrifuged for 45 min at 50,000 x *g* at 4°C in a Sorvall table-top ultracentrifuge with a TLA 100 fixed angle rotor. The supernatants were collected, and the pellets were resuspended in RNA granule buffer. The proteins were precipitated overnight at -20°C by adding 7 ml of chilled 100% ethanol to the collected supernatant and pellet samples. The precipitated samples were then centrifuged for 45 min at 2177 x *g* at 4°C in an Eppendorf 5810 swing bucket rotor and protein pellets were resuspended in 1x SDS sample buffer diluted in RNA granule buffer and separated by SDS-PAGE.

### Nuclease Treatment and separation of monosomes for immunoblots

We digested the GF into monosomes to see whether the puromycylated peptides remained on individual ribosomes as previously described (17). 100 U of RNaseI [100 U/µl; Ambion AM2294, Thermo Fisher] was added to the puromycylated GF and incubated for 30 minutes at 4°C on a rocker. The nuclease was quenched with 100 U of SuperaseIN [20 U/µl; Invitrogen #AM2696, Thermo Fisher], and the samples were loaded onto a fresh 15-60% sucrose gradient and centrifuged for 90 minutes at 56660 x *g* at 4°C in an AH-629 swing bucket rotor to separate monosomes. In some cases, nuclease was omitted to test if ribosomal subunits were dissociated by puromycin. Fractions of 3.5 ml were collected from the top, and the pellet fraction was resuspended in RNA granule buffer. The fractions were precipitated overnight at -20°C by adding 7 ml of chilled 100% ethanol to the collected supernatant and resuspended pellet samples. The precipitated samples were then centrifuged for 45 min at 2177 x *g* at 4°C in an Eppendorf 5810 swing bucket rotor and protein pellets were resuspended in 1x SDS sample buffer diluted in RNA granule buffer and separated by SDS-PAGE.

### Immunoblotting

Samples were run on 12% polyacrylamide gels, transferred onto 0.45µm nitrocellulose membranes [Bio Rad; 1620115] by wet transfer and blocked with 5% bovine serum albumin (BSA) [Sigma; A9647] in 1X Tris-Buffered Saline with Tween (TBS-T). Membranes were incubated with primary antibodies (anti-puromycin AB 2619605 Developmental Studies Hybridoma Bank) (1:1000) for 1 hour at room temperature, followed by HRP-conjugated secondary antibodies [ThermoFisher; #31430, #31460] and detected by chemiluminescence [Perkin Elmer; NEL105001EA]. In some cases, the blots were then stripped [ZmTech Scientific; S208070] and reblotted with an anti-eS6 antibody [Cell Signaling #2217] or with anti-L7 (uL20) antibody [Proteintech 14583-1-AP]. Signal was quantified using ChemiDoc Imager software.

### Isolation of monosomes for Cryo-EM

For cryo-EM experiments, fractions 2 and 3 containing monosomes collected from the sucrose gradient were diluted with RNA granule buffer and loaded onto a 2 ml 20% sucrose [Calbiochem; 8550] cushion (dissolved in RNA granule buffer) in a Sorvall 36 ml tube [Kendro; 3141, Thermo Scientific], filled to top with additional RNA granule buffer and centrifuged for 2 hours in a Thermo Scientific AH-629 swing-bucket rotor at 56660 x *g* at 4°C to achieve a monosome pellet. The monosome pellet was resuspended in 100 µL of RNA granule buffer and loaded onto grids for cryo-EM (See Cryo-electron microscopy). Alternatively, after treating the GF with puromycin and nuclease, samples were directly applied to cryo-EM grids.

### Cryo-electron microscopy

For all cryo-EM samples a volume of 3.6 μL of monosome solution at a concentration of 170 nM (for all cryo-EM samples) was applied to holey carbon grids (C-flat CF-2/2–2Cu-T) that had been glow discharged in air at 10 mA for 18 seconds. Grids were blotted in a Vitrobot Mark IV (Thermo Fisher Scientific Inc.) for 3 seconds with a blot force of +1 before plunging into liquid ethane. The Vitrobot chamber was set to 25 °C and 100% relative humidity.

Cryo-EM images were collected using SerialEM software (44) in the Titan Krios at FEMR-McGill (**Table 1**). Movies were recorded in a Gatan K3 direct electron detector with a Quantum LS imaging filter. The total dose used for each movie was 50 e/Å^2^ equally spread over 30 frames. Images were collected at a magnification of 81,000x, yielding images with a calibrated pixel size of 1.09 Å. The nominal defocus range used during data collection was between -1.25 and -2.75 μm.

**Table 1.** Cryo-EM analysis of puromycylated 80S ribosomes from RNA granules. Data acquisition parameters and map statistics.

|  | 80S Class 1<br>(tRNA A/P & P/E) | 80S Class 2<br>(tRNA P/P) | 80S Class 3<br>(Puromycin bound) | Free 60S Class 3 |
| --- | --- | --- | --- | --- |
| Data collection |  |  |  |  |
| Microscope | Titan Krios |  |  |  |
| Detector | Gatan BioQuantum LS K3 |  |  |  |
| Nominal Magnification | 81,000x |  |  |  |
| Voltage (kV) | 300 |  |  |  |
| Total exposure (e-/Å²) | 50 |  |  |  |
| Number of frames | 30 |  |  |  |
| Defocus range (µm) | -1.25 to -2.75 |  |  |  |
| Calibrated physical pixel size (Å/px) | 1.09 |  |  |  |
| Reconstruction and refinement |  |  |  |  |
| Particle number | 485,712 | 86,464 | 405,839 | 463,036 |
| Resolution (Å) | CM: 3.1 Å<br>MBR: 60S (3 Å) | CM: 3.3 Å<br>MBR: 60S (3.2 Å) | CM: 3.1 Å<br>MBR: 60S (3.1 Å) | CM: 3.1 Å |
| Data Deposition |  |  |  |  |
| EMDB code | EMDB-40665 | EMDB-40666 | EMDB-40667 | EMDB-40668 |

CM: Consensus map; MBR: Multi-body refinement.

### Image processing

Cryo-EM movies were corrected for beam-induced motion using RELION’s implementation of the MotionCor2 algorithm (45, 46). We used 5 x 5 patches, no frame grouping, B factor 150 and dose-weighting (Supplementary Figure 2). Only dose-weighted averages were saved. CTF parameter estimation was done using CTFFIND-4.1 program (47) using the dose-weighted averages with a 30-5.0 Å resolution range and 512-pixel FFT box size. Minimum and maximum defocus values were set up at 1,000 and 50,000 Å, and the defocus step size was set at 100 Å. Only micrographs with a resolution estimated to 8 Å or better were selected for further processing. The remaining processing steps were done using RELION-4.0, and the workflow is shown in Supplementary Figure 2 (46). Particles were automatically picked by template matching. We first picked 18,802 particles from 129 micrographs collected at various defocuses using the Laplacian picker to obtain the templates for particle auto-picking. The minimum and maximum diameter for the blob-detection algorithm was 200 and 434 Å, respectively. The maximum resolution considered in the micrographs was 20 Å, and the default and upper threshold were set to 0.4 and 4. These particles were subjected to one round of reference-free 2D classification requesting 50 classes to generate the templates for auto-picking in the dataset comprising 13,716 movies. In this process, templates were low pass filtered at 20 Å, and we used a picking threshold of 0.2 and a minimum inter-particle distance of 120 Å. The 3,490,854 particles selected were extracted from the dose-weighted summed micrographs, further binned to a 5.70 Å/pixel and 76-pixel box size and then subjected to two rounds of 2D classification for the particle curation process. We requested 150 classes in the first round of 2D classification and 100 classes in the second round. We used a regularization parameter (tau fudge) value of T = 2 for all 2D classifications. The particles from the best-aligned classes generated a particle stack of 1,664,649 particles. To separate the particles representing the various types of ribosomes present in the sample, we first performed two layers of exhaustive 3D classification, requesting 5 classes in the first layer and 5 classes per class in the second layer (Supplementary Figure 3; top panel). All 3D classifications used a regularization parameter value of T = 4, ran for 25 iterations and used a circular mask of 434 Å. The initial 3D reference used for the exhaustive 3D classification was an 80S neuronal ribosome map from rats low-pass filtered at 60 Å created from EMD-28726 and EMD-28727 (17). The resulting maps from the exhaustive 3D classification were visually inspected in Chimera (48) and it was concluded that there were two main classes of particles. The 2D projections assigned to each class were pooled and subjected to a 3D auto-refine process using a circular mask of 434 Å to generate the consensus refinement maps for the two structures. The particle orientation parameters from these consensus structures were used in a 3D classification step without image alignment (Supplementary Figure 3; middle panel). In this process, two additional classification layers were done, generating four different classes. Particles corresponding to each class were again sent to 3D auto-refine to produce consensus maps. To finally separate the particles in the dataset according to their features in the functional site, the four consensus structures were used as references for an additional multi-layer 3D focused classification process with each of the classes using a spheric soft-mask (4-pixel extension, 6-pixel soft cosine edge) around the A, P and E sites (Supplementary Figure 3; bottom panel). Those particles representing similar structures in this region were pooled together, obtaining a final number of four classes. To remove any ‘contaminating’ 60S free subunits particles potentially included in the three classes showing entire 80S ribosomes, each of these classes was subjected to an additional focused classification step using a dilated mask encompassing the entire 40S. We found no evidence for this type of particle misclassification. High-resolution refinements of the four obtained classes were performed in five stages: In the first step, particles from each class were re-extracted with the original pixel size (1.09 Å/pixel, 398-pixel box size) and subjected to a 3D auto-refine process with a 434 Å circular mask and using as an initial model the maps obtained via 3D classification (after proper scaling and filtering using a 60 Å low-pass Fourier filter). The resulting maps were used as the initial model for a second refinement step. This second step of 3D auto-refine processing used a tight mask created from the maps obtained in the first refinement step. The binarization threshold used to create this mask was selected using Chimera (48), and we also extended the binary mask by 4 pixels and added a 4-pixel soft cosine edge. The outputs of the second 3D auto-refine and subsequent postprocessing processes were used in the third refinement step involving CTF refinement with ‘Estimate (anisotropic) magnification’ set as “yes” in the first cycle. In a second cycle of CTF refinement, we selected ‘Perform CTF parameter fitting’ as “Yes” and selected ‘Fit defocus’ as ‘Per-particle’ and ‘Fit astigmatism as ‘Per-micrograph.’ We also selected ‘No’ for ‘Fit B-factor’, ‘Fit phase-shift’, ‘Estimate trefoil’ and ‘Estimate 4th order aberrations. However, we selected ‘Yes’ for ‘Estimate beam tilt, as our data was collected using this approach. We also selected ‘Yes’ for ‘Estimate trefoil’ and ‘Estimate 4th order aberrations’. In the fourth refinement stage, we used the particles of the CTF refinement process to run Bayesian polishing to correct for per-particle beam-induced motion before subjecting these particles to a final round of 3D refinement to generate high-resolution maps (fifth refinement step). Bayesian polishing was performed using sigma values of 0.2, 5,000 and 2 for velocity, divergence and acceleration, respectively. To better define the density of the 40S subunit for the structure of each one of the classes, we used the particles assigned to each class and performed multi-body refinement by dividing the consensus cryo-EM map into two bodies (body 1: 60S and body 2: 40S plus tRNA). A soft mask was generated for each body and applied to the corresponding map during refinement. Sharpening of the final cryo-EM maps and average resolution estimation, and local resolution analysis were done with RELION using the gold-standard approach (49, 50). Cryo-EM map visualization was performed in UCSF Chimera (48) and Chimera X (51). The structures are available on the EMDB (https://www.ebi.ac.uk/emdb/ ; accession codes in Table 1).

The second dataset obtained from puromycin-treated monosomes purified as for Anadolu et al. (17) was analyzed using an identical image processing pipeline. In this case, we collected 13,645 movies and use a subset of 207 movies to initially pick up 19,733 particles using the Laplacian picker and generate the templates for particle picking in the entire dataset. A total of 1,797,434 particles were selected from all the movies. Particle curation using 2D classification generated a total of 445, 607 particles that were subjected to 3D classification.

To obtain the 2D class averages from the particles in each class (Supplementary Figure 5) the 2D classification jobs for each group of particles using a regularization parameter value of T = 2.

To explore the movement of the 40S subunit in classes 1 and 2 in the -puromycin sample and classes 1, 2 and 3 in the +puromycin sample, we used 3D variability analysis as implemented in CryoSPARC v4 (52, 53). Each set of particles was analyzed separately. Particles belonging to each set were re-extracted, binned to a pixel size of 4.36 Å/pixel and 100-pixel box size. Each particle stack was subjected to one cycle of ab-initio 3D reconstruction. For this step, we selected 0.85 and 0.99 as inner and outer window radius, requested 3 classes and a maximum and minimum resolution to consider of 35 Å and 12 Å. The number of iterations before and after annealing starts and ends was set to 200 and 300, respectively and the increase of Fourier radius at each iteration was 0.04. All other parameters for this routine were used with the default settings and values. The best particles and model from the ab-initio 3D reconstruction process were selected for subsequent Non-Uniform Refinement, which was run under default settings with C1 symmetry, optimized per-particle defocus, optimized per-exposure group CTF parameters and options ‘Fit Spherical Aberration’, ‘Fit tetrafoil’ and ‘Fit anisotropic Magnification’ deactivated. The aligned particles were used for 3D variability analysis requesting 3 orthogonal principal modes. A binary mask with dilation radius 4 (pix) and soft padding width 15 (pix) focusing on the 40S subunit was applied during analysis, and the result was displayed in simple mode with 8 frames and with all settings at default values. Results were filtered at 10 Å resolution. Results were visualized in Chimera (48, 51) by assembling movies from the obtained frames.

### Cultures used

Rat primary hippocampal neurons and cortical neurons were dissected from embryonic day 18 Sprague-Dawley embryos and cultured as previously described (54) (16). HEK293T cells were cultured in DMEM (Life Technologies, Burlington, Ontario) supplemented with 10% fetal bovine serum, sodium pyruvate, penicillin and streptomycin.

### Ribopuromycylation of cell cultures

Ribopuromycylation was performed as described (13) in 8-10 DIV primary rat hippocampal neurons using 0.0003% digitonin for the extraction step. Cells were imaged using a Zeiss LSM-710 confocal microscope with a 63X oil immersion objective (NA=1.4). ImageJ was used for image post-processing, including straightening of neurites (using the “Straighten Curved Objects plugin”) and quantitation. For quantitation of puromycylated puncta, straightened images of neurites were thresholded so that only high intensity puncta were visible, and their numbers were counted and normalized to neurite length.

### Inhibition of Puromycylation by Anisomycin

Rat hippocampal neurons and cortical neurons were incubated for five minutes in cell solution (neurobasal containing N2, B27, glutamax and penicillin) containing either no puromycin, 100 µM puromycin, or 100µM puromycin and 100 µM anisomycin. HEK293T cells were incubated in the same concentration of puromycin or puromycin and anisomycin for 5 minutes in DMEM + 5% FBS cell solution. The hippocampal neurons, cortical neurons, and HEK293T cells were washed, then lysed on ice for 15 minutes in non-denaturing lysis buffer (150 mM KCl, 10 mM MgCl_2_, 5mM HEPES, 1% IGEPAL) supplemented with protease inhibitors (Roche Complete EDTA-free, Roche, Laval, Canada). The samples containing lysis buffer were then collected and spun down for 5 minutes in pellet the cell debris. The supernatant was added to the SDS sample buffer for immunoblotting before loading onto a 12% acrylamide gel. The resolved proteins were transferred onto a 0.45 μm nitrocellulose membrane [Bio Rad; 1620115] for Ponceau staining ([Ponceau S: BP103-10, FisherBiotech]) and immunoblotting. The membranes were blocked with 5% BSA [Sigma; 9647] in Tris Buffered Saline with Tween (TBS-T [Tris: Fisher, BP152-1; NaCl: Fisher, BP358-212; Tween: Fisher, BP337]) before incubation with anti-puromycin (see reagents) at 1:1000 to 1:2000 dilution. Membranes were washed with TBS-T after incubation. Detection was done using HRP-conjugated secondary antibodies [ThermoFisher; #31430, #31460] followed by ECL [Perkin Elmer; NEL105001EA] reaction and imaging using the Bio Rad ChemiDoc digital imager. Quantification of signal intensity was done using ImageJ software. We selected full lane ROIs and quantified staining in the entire lane. We similarly quantified images of the Ponceau stained membrane before immunoblotting. An anti-puromycin/Ponceau ratio was calculated for each lane. The background in the absence of puromycin was then subtracted from these values. The resulting value in the presence of anisomycin was then divided by the value in its absence to detect the resistance to anisomycin which varied between 1 (completely resistant) to 0 (completely sensitive).

### Statistical analysis

ANOVA with post-hoc Bonferroni tests was used to detect differences in the competition between anisomycin and puromycin.

## Supporting information

Vdieo 2

## Acknowledgements

We thank the Facility for Electron Microscopy Research (FEMR) staff at McGill University. FEMR is supported by the Canadian Foundation for Innovation, the Quebec government, and McGill University. We thank Taylor Stephens for assistance with Supplemental Figure 5.

## Data Availability

The cryo-EM maps obtained in this study have been deposited in the Electron Microscopy Data Bank (EMDB), and the accession codes are detailed in Table 1.

## Supplementary Figure Legends

**Supplementary Figure 1.**
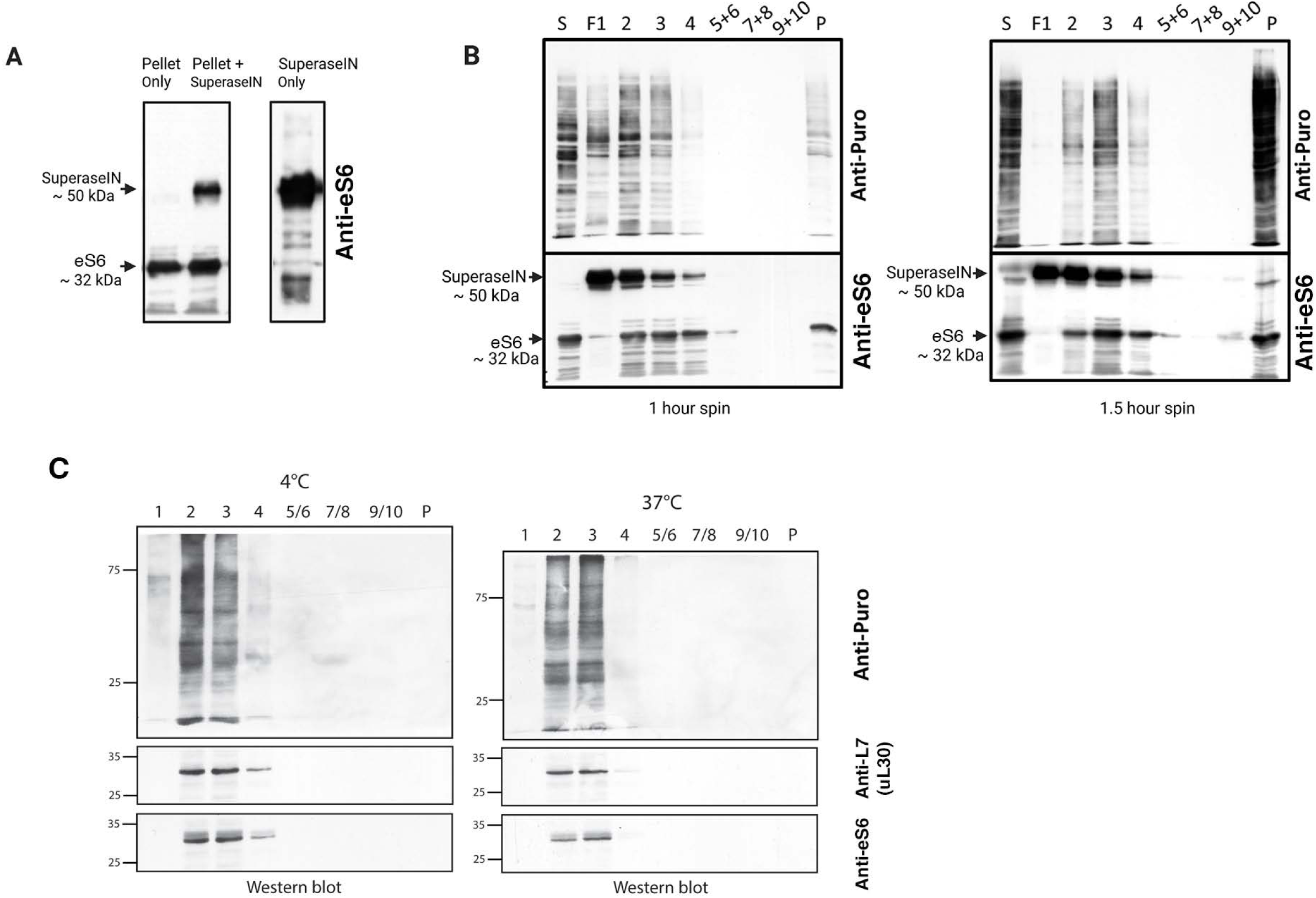
Additional data on biochemical fractionation. **A)** Anti-eS6 immunoblots of samples from Pellet, Pellet +SuperaseIN and SuperaseIN alone. The ∼50 kDa band was only observed when SuperaseIN was added (left panel). When SuperaseIN was run alone on an SDS gel, the same sized band was observed, confirming that the anti-S6 antibody has cross-reactivity with SuperaseIN (right panel). B) Replicates of the experiment displayed in Fig. 1D; immunoblots of sucrose gradient fractions after nuclease treatment, blotted with anti-puromycin (top) and anti-eS6 (bottom) antibodies. In the left panel, the centrifugation was only 1 hour, yielding some residual ribosomes in F1 soluble fraction. Whereas, when the spin time was increased to 1.5 hours, we observed better separation between soluble fraction F1 and monosomal fraction F2. The same 1.5hr spin was used in Fig. 1D. Note the incomplete nuclease digestion of the GF (P) in example 2 led to a large increase in S6 and puromycylated proteins in the pellet fraction. Lane labelled with S represents the puromycylated GF before fractionation. C) Comparison between fractionation of puromycylated peptides when puromycylation was at 4^0^C or at 37^0^C. In this example, the starting material (S) was not loaded. Anti-L7 was used to monitor the large subunit and the anti-eS6 is also shown for comparison. This experiment was performed two additional times with similar results.

**Supplementary Figure 2.**
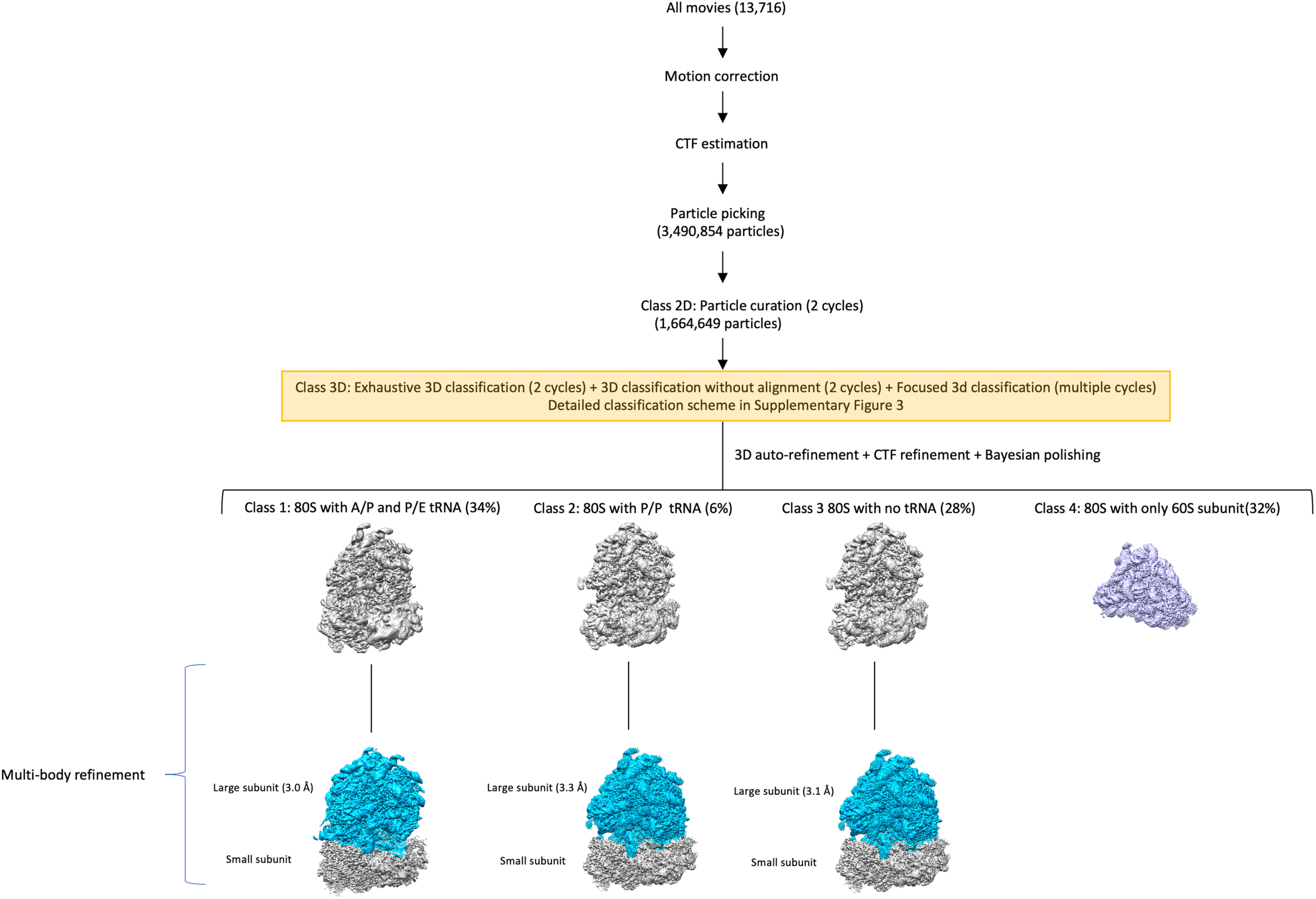
Cryo-EM analysis processing workflow of puromycin-treated stalled neuronal ribosomes. Image processing workflow of the cryo-EM datasets collected from the 80S ribosomes in the granule fraction of the sucrose gradient after nuclease digestion and puromycin treatment. The diagram displays the main image processing steps undertaken with these datasets and the four main ribosome populations found. The class 3D step is highlighted in yellow to indicate that the detailed protocol of the classification strategy is shown in Supplementary Figure 3. The number of movies and particle images maintained at each step is indicated. The resolutions of the cryo-EM maps obtained for the large subunit through multibody refinement are also indicated. The maps shown in the figure were generated from the dataset puromycin-treated monosome sample purified according to the protocol in Figure 1A & C.

**Supplementary Figure 3.**
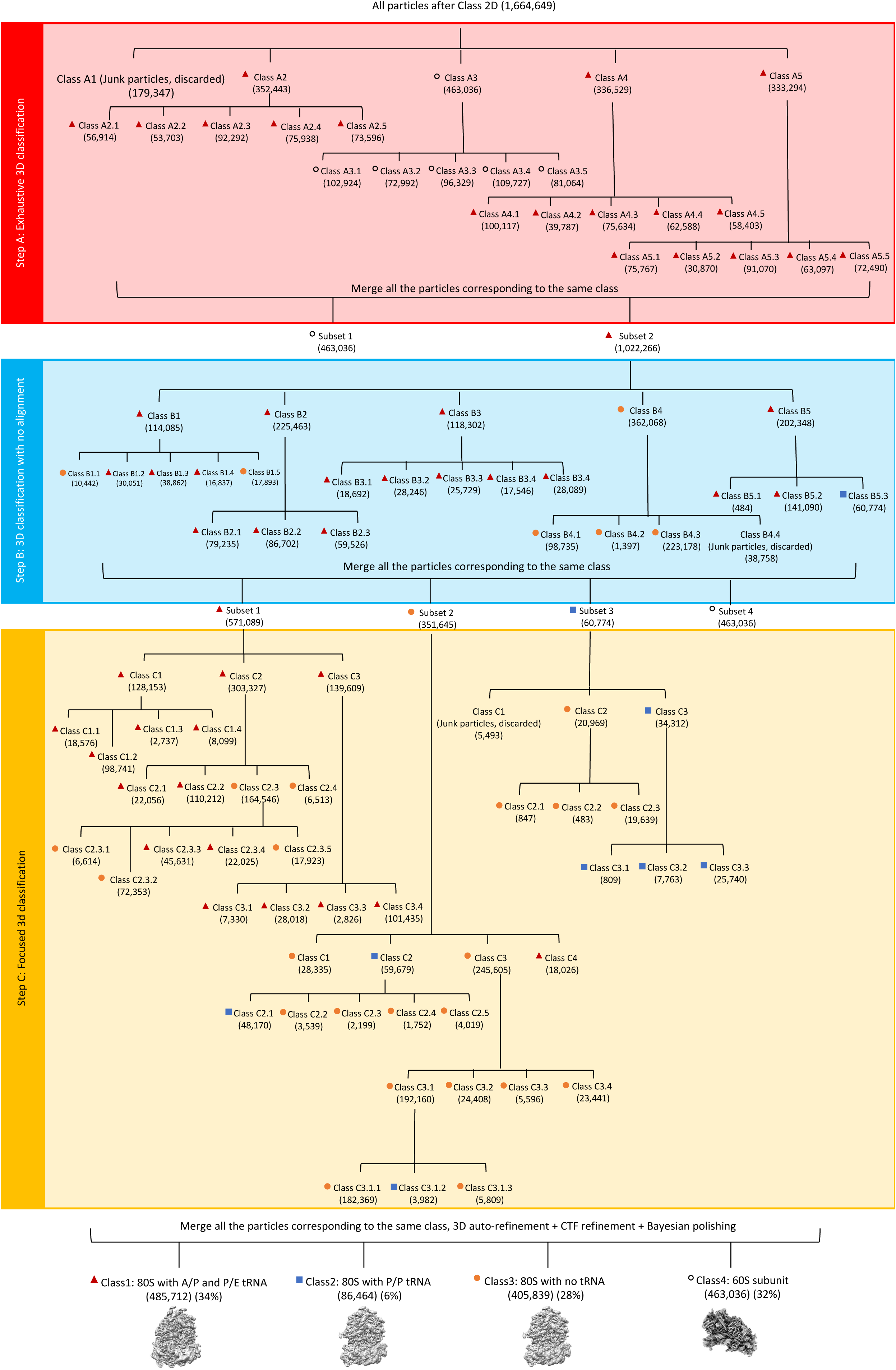
3D classification scheme for the puromycin-treated stalled neuronal ribosomes. The diagram shows the detailed classification scheme for the three steps of classification. Each step is highlighted in a different color. The diagram shows the number of classes obtained at each step. The number of particles assigned for each class is indicated by the number in brackets. As the classification progresses, each group of particles is assigned to one of the final classes, and this is indicated with a red triangle (class 1), blue square (class 2), orange circle (class 3) or white circle (class 4) next to the name of the class.

**Supplementary Figure 4.**
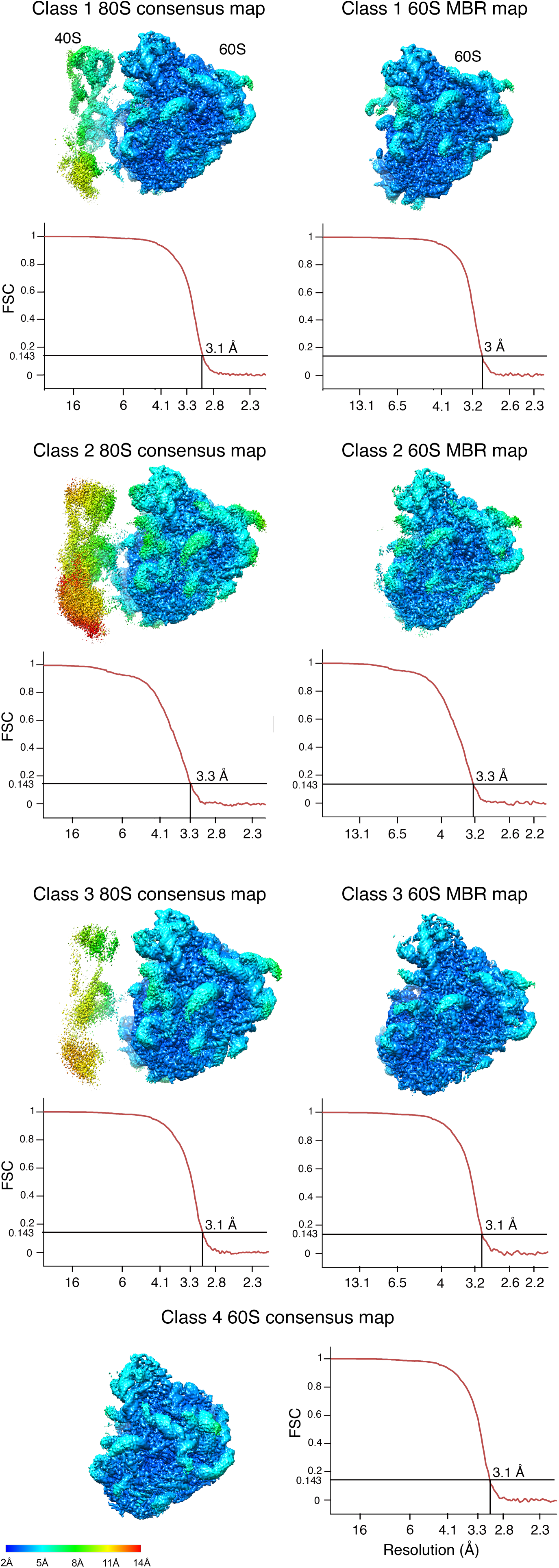
Resolution analysis of the cryo-EM maps obtained for the puromycin-treated stalled neuronal ribosomes. Particles for each class were refined into a consensus map. These maps were subsequently refined by multi-body refinement (MBR) by dividing the 80S ribosome into two major bodies, the 40S and the 60S particles. The Fourier shell correlation graphs for the consensus maps and the 60S subunits after the maps were subjected to MBR are shown for all classes. We used a FSC threshold of 0.143 to report the resolution. Maps are colored according to their local resolution using the color coding indicated in the scale bars. Ribosomal subunits are indicated in class 1.

**Supplementary Figure 5.**
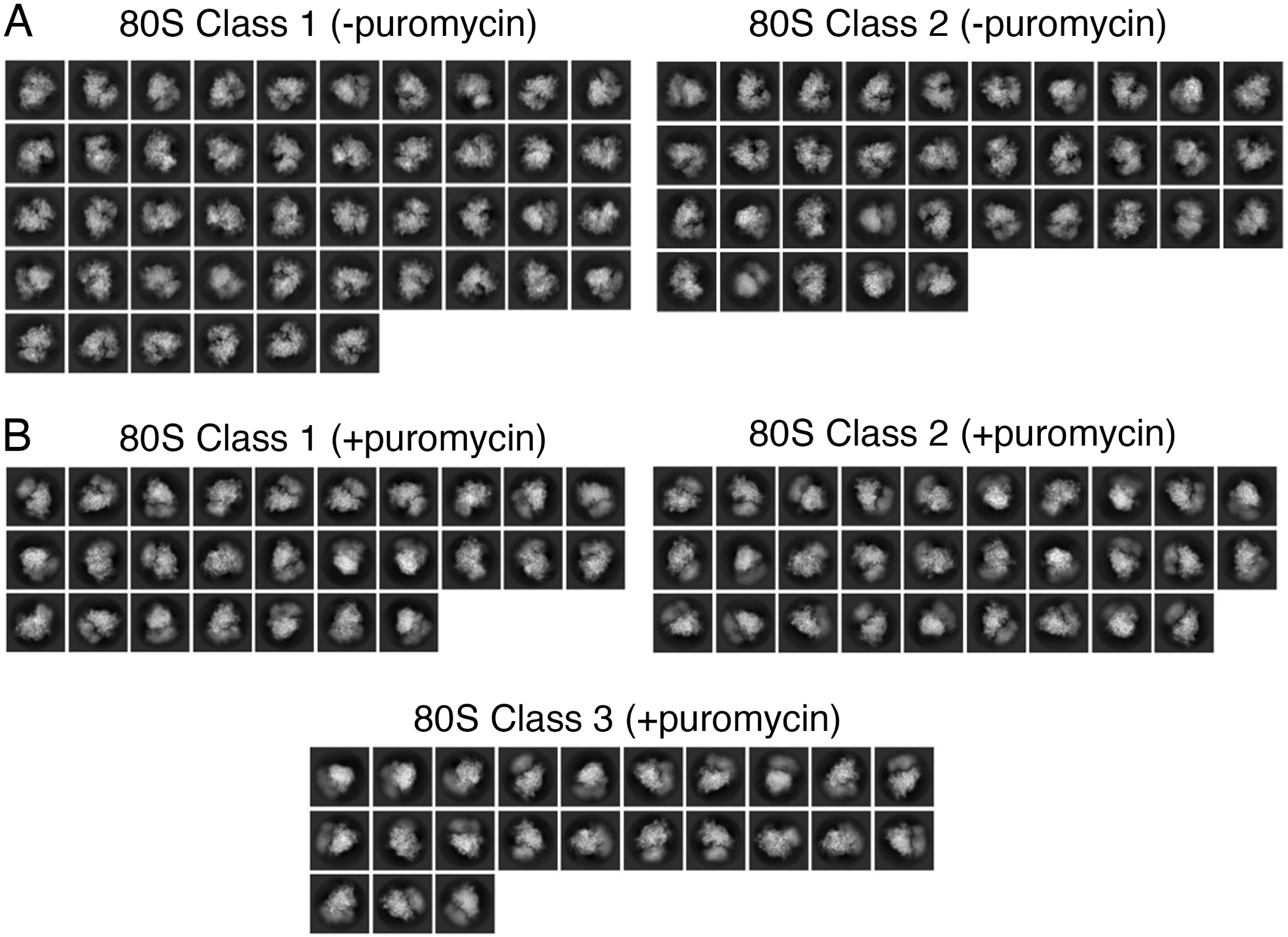
2D class averages for all classes obtained in the untreated and puromycin-treated samples. Each panel shows a group of 2D class averages obtained from particles assigned to each class obtained for the untreated (A) and puromycin-treated (B) samples.

**Supplementary Figure 6.**
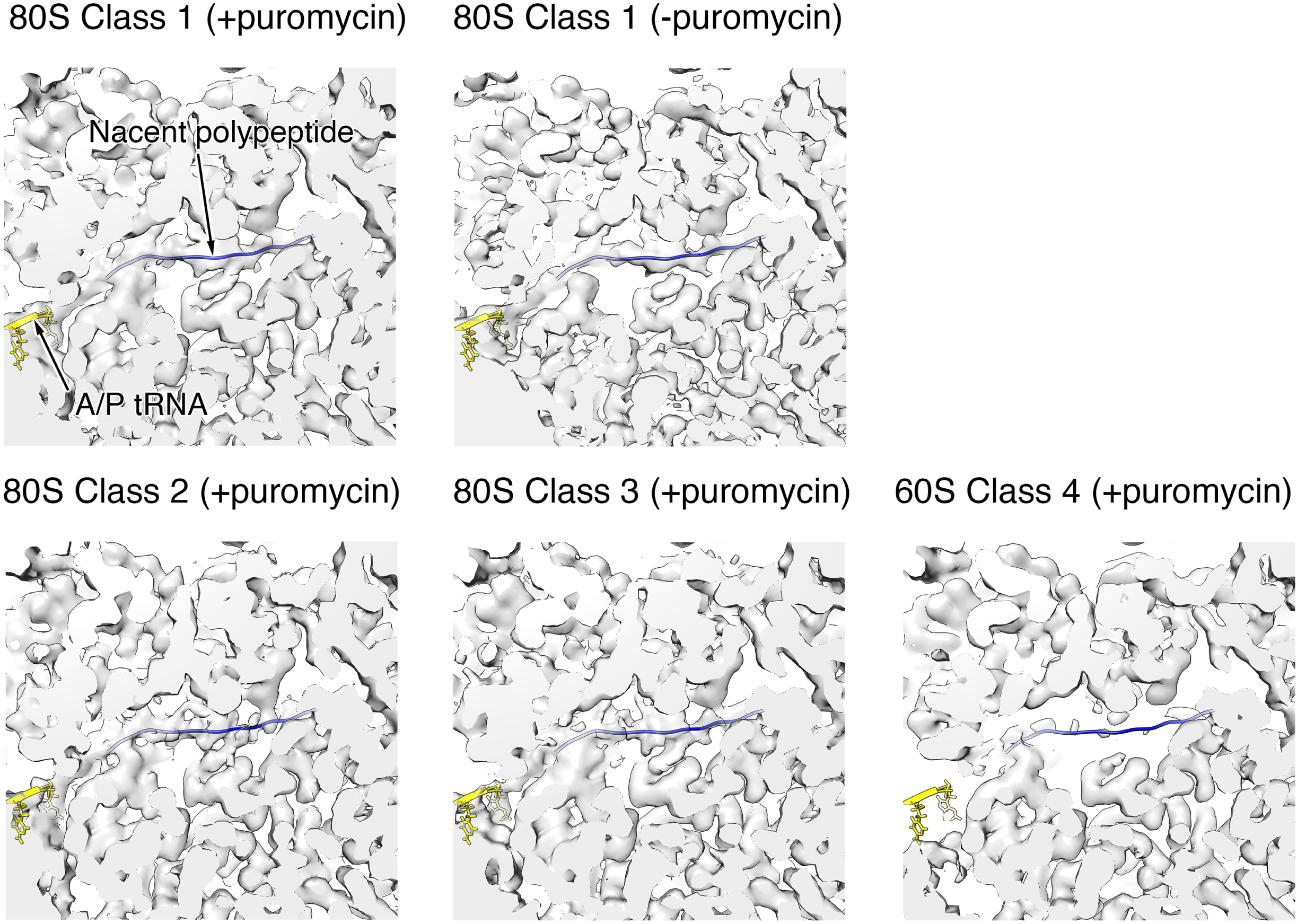
Nascent chain polypeptides are retained in the puromycin treated ribosomes. The central section of the 80S ribosomes class 1 to 4 in the puromycin-treated sample shows the nascent chain’s path. The top middle panel also shows a comparative view of the same area in the 80S ribosome from class 1 in the untreated sample. The CCA 3’end of the A/P tRNA from pdb 7NWG is shown in yellow. The path of the nascent peptide is shown by a poly-alanine chain from pdb 6HCJ colored in blue and docked into the density.

**">Supplemental Figure 7.**
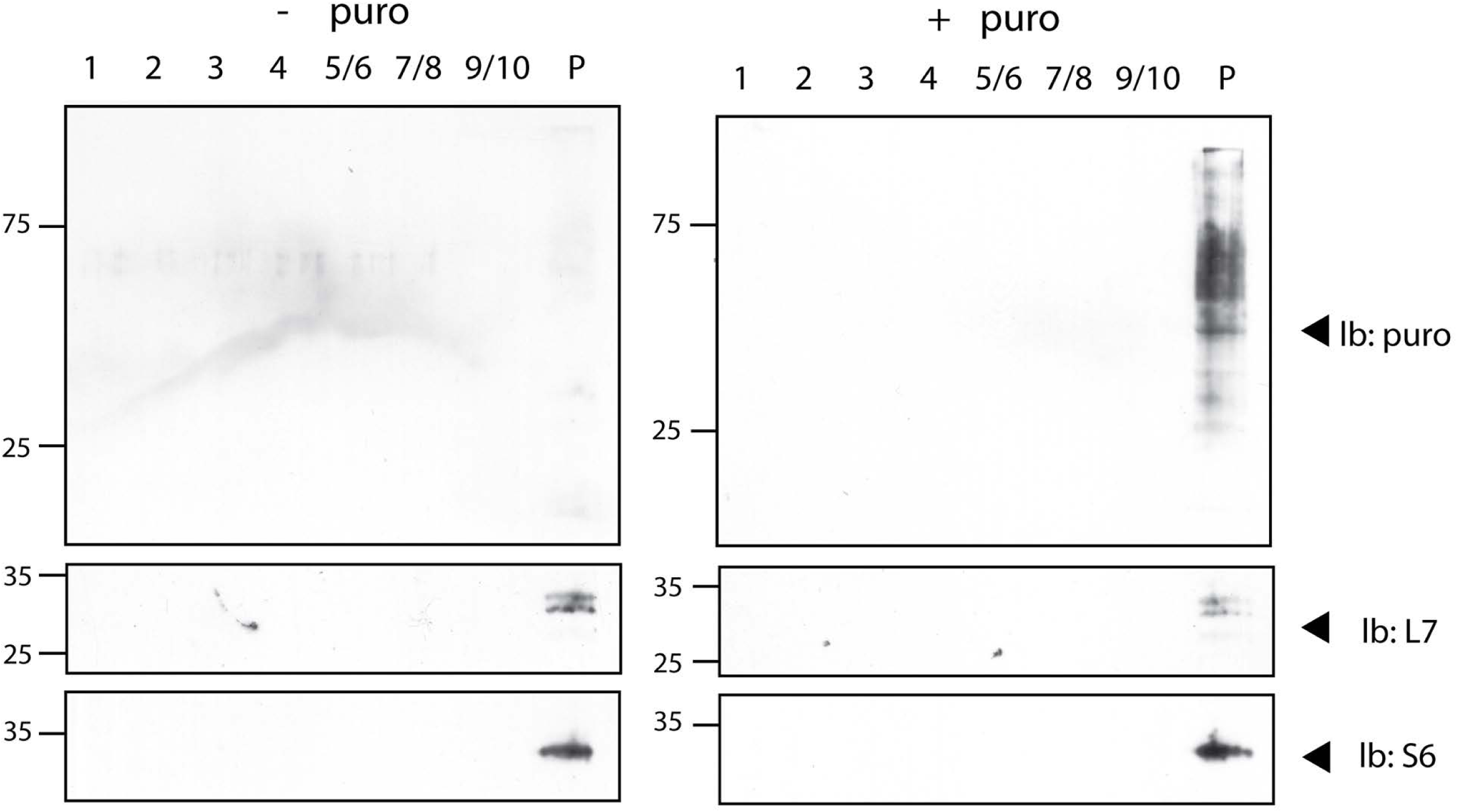
No dissociation of ribosomal subunits is observed after puromycylation. Immunoblots with anti-puromycin, anti-eS6 and anti-L7 of resedimented granule fractions either in the presence or absence of puromycylation. No puromycylated peptides or ribosomal subunit staining is observed in fractions corresponding to dissociated ribosomal subunits. This experiment has been repeated three times and no dissociated ribosomal subunits were detected in any of the experiments.

**Supplementary Figure 8.**
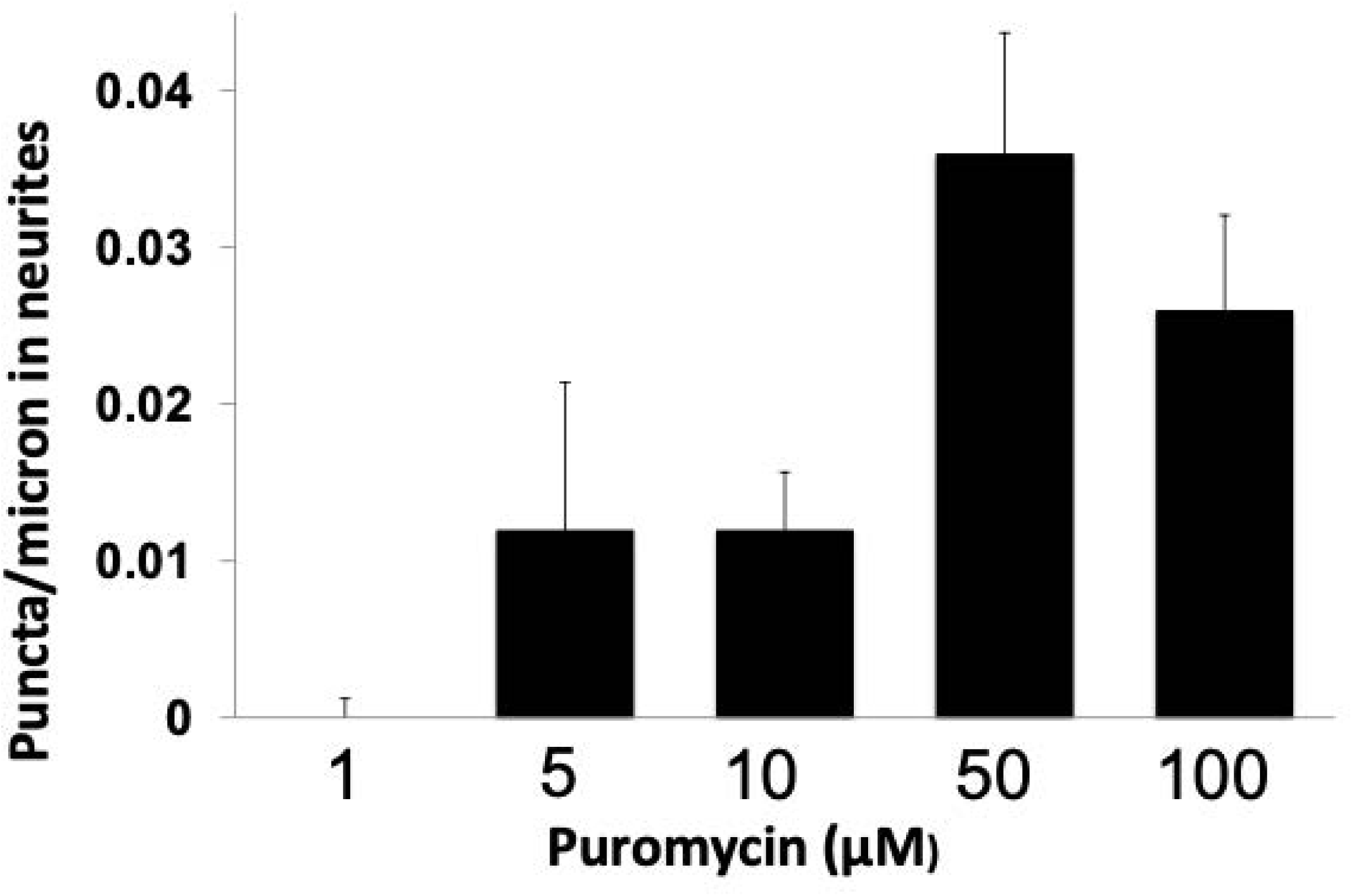
Ribopuromycylation of stalled polysomes requires higher concentrations of puromycin in neurons. The density of homoharringtonine resistant ribopuromycylation puncta in hippocampal dendrites was measured at five different concentrations of puromycin. The values represent the average +/-S.E.M. from three independent cultures, with a minimum of 12 neurites measured in each culture.

## Videos

**Video 1.** Video showing the 40S detachment motion in class 1 (-puromycin): 80S with A/P and P/E tRNA.

**Video 2.** Video showing the 40S detachment motion in class 2 (-puromycin): 80S with P/P tRNA.

**Video 3.** Video showing the ratcheting motion in class 2 (-puromycin): 80S with P/P tRNA.

**Video 4.** Video showing the 40S detachment motion in class 1 (+puromycin): 80S with A/P and P/E tRNA.

**Video 5.** Video showing the ratcheting motion in class 1 (+puromycin): 80S with A/P and P/E tRNA.

**Video 6.** Video showing the 40S sliding motion in class 1 (+puromycin): 80S with A/P and P/E tRNA.

**Video 7.** Video showing the 40S detachment motion in class 2 (+puromycin): 80S with P/P tRNA.

**Video 8.** Video showing the ratcheting motion in class 2 (+puromycin): 80S with P/P tRNA.

**Video 9.** Video showing the 40S sliding motion in class 2 (+puromycin): 80S with P/P tRNA.

**Video 10.** Video showing the 40S detachment motion in class 3 (+puromycin): 80S with no tRNA.

**Video 11.** Video showing the ratcheting motion in class 3 (+puromycin): 80S with no tRNA.

**Video 12.** Video showing the 40S sliding motion in class 3 (+puromycin): 80S with no tRNA.

